# Comparative transcriptome reprogramming in oak galls containing asexual or sexual generations of gall wasps

**DOI:** 10.1101/2024.04.09.588582

**Authors:** Eleanor Bellows, Matthew Heatley, Nirja Shah, Nathan Archer, Tom Giles, Rupert Fray

## Abstract

- Oak gall wasps have evolved strategies to manipulate the developmental pathways of their host to induce gall formation. This provides shelter and nutrients for the developing larva. Galls are entirely host tissue; however, the initiation, development, and physical appearance are controlled by the inducer. The underlying molecular mechanisms of gall formation, by which one or a small number of cells are reprogrammed and commit to a novel developmental path, are poorly understood. In this study, we sought a deeper insight into the molecular underpinnings of this process.
- Oak gall wasps have two generations each year, one sexual, and one asexual. Galls formed by these two generations exhibit a markedly different appearance. We sequenced transcriptomes of both the asexual and sexual generations of *Neuroterus quercusbaccarum* and *Neuroterus numismalis*. We then deployed Nanopore sequencing to generate long-read sequences to test the hypothesis that gall wasps introduce DNA insertions to determine gall development.
- We detected potential genome rearrangements, but did not uncover any non-host DNA insertions. Transcriptome analysis revealed that the transcriptomes of the sexual generations of distinct species of wasp are more similar than inter-generational comparisons from the same species of wasp.
- Our results highlight the intricate interplay between the host leaves and gall development, suggesting that season and requirements of the gall structure play a larger role than species in controlling gall development and structure.

**Summary Statement:** Oak gall wasps, *Neuroterus quercusbaccarum* and *Neuroterus numismalis*, induce species-specific galls on *Quercus robur* leaves. We demonstrate that the sexual generation of distinct species of wasps induce more similar changes in the host than different generation galls from the same species.

## 1. Introduction

Plant galls are structures induced by parasites that provide shelter and nourishment to the inducer. Oak galls are predominantly caused by gall wasps of the family Cynipidae and can be structurally complex. These small wasps lay their eggs on different tissues of the oak tree - inducing an oak gall. In the UK, more than 50 different types of oak gall can be found. The different species of oak gall wasps can be identified by the type of gall that they induce. Thus, despite being entirely made up of plant-derived cells, the initiation, development, and physical appearance is determined by the insect. This phenomenon makes oak galls a classic example of an extended phenotype (Dawkins and Dennett, 1999).

The asexual and sexual generations of *Neuroterus quercusbaccarum* and *Neuroterus numismalis* lay their eggs in the underside of *Q. robur* leaves; the asexual generation of *N. numismalis* can also lays its eggs in the catkins of the tree. Galls of the same generation but different species can be found in close proximity to each other on the same leaves. These oak gall species are rare examples of cyclical parthenogenesis (or heterogony). Each year, there is one asexual generation and one sexual generation that always follow each other (Stone et al., 2002). In spring the sexual generation of *N. quercusbaccarum* and *N. numismalis* emerge and lay their eggs, producing currant and blister galls, respectively, each gall contains a single larva (Figure 1a-c). The oviposition initiates the growth of the gall, which grows to house the developing larva (Stone et al., 2002). The asexual generations emerge in early summer and lay their eggs, producing rough spangle galls and silk button galls respectively, again each containing a single larva (Figure 1d-f). These galls drop off the leaves in autumn, and the wasp hibernates inside until spring, when it emerges to start the cycle again. It is hypothesised that oviposition site and timing are very important for gall formation and that meristematic tissue is needed for gall initiation (Stone et al., 2002). It has been noted that the eggs are always laid on young leaves, the appearance of the different generations occurs at the same times as the first and second flushes of leaves in spring and early summer (Hough, 1953).

**Figure 1.**
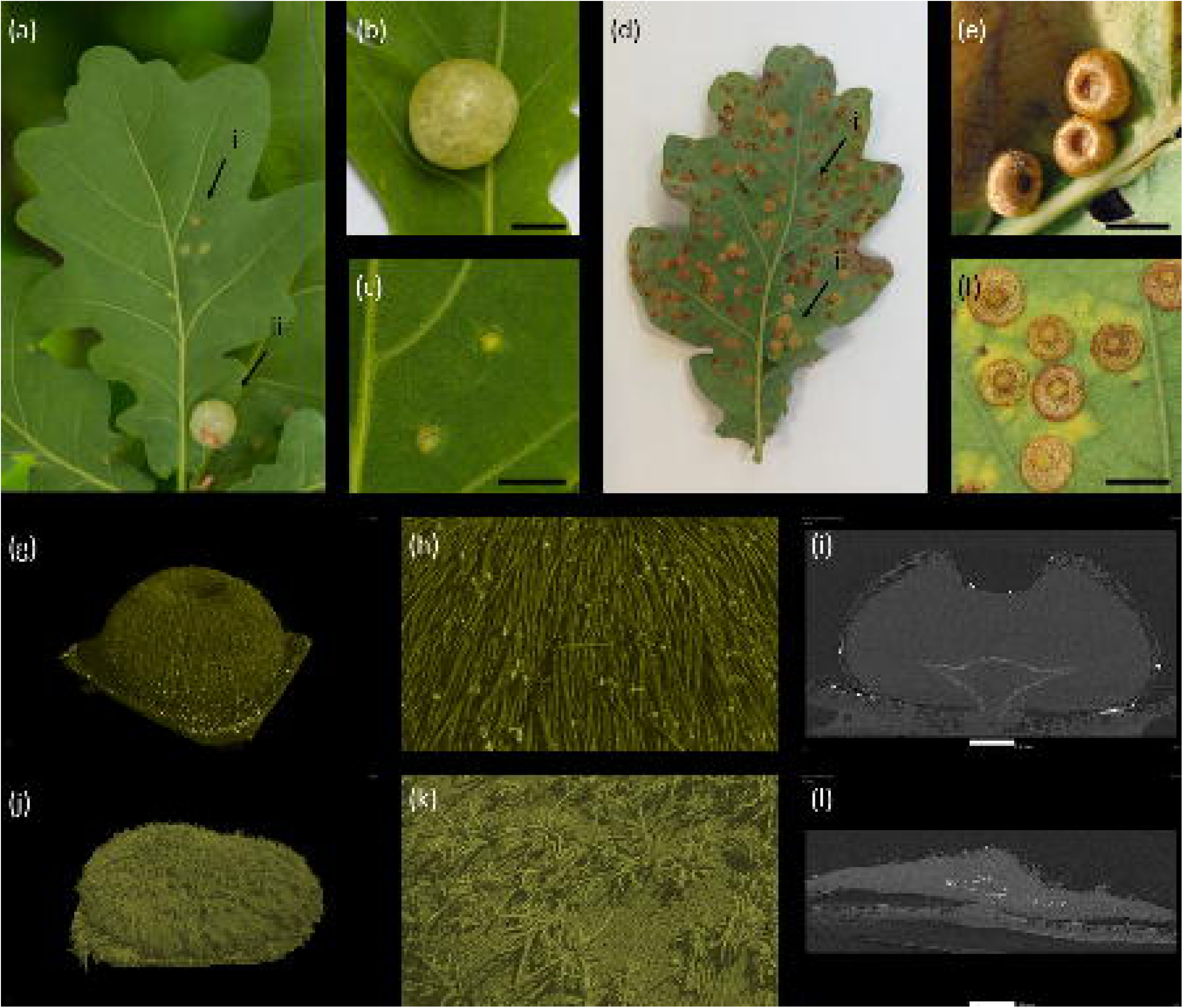
Representative images and CT scans of the galls used in this study. (a) the galls containing the asexual generation of the *Neuroterus numismalis*and *Neuropterus quercusbaccarum* species, (i) blister and (ii) currant galls respectively, appear at the same time of year on the same leaves. (b) currant gall on an oak leaf, scale bar is 1 cm. (c) blister gall in an oak leaf, scale bar is 1 cm (d) the galls containing the sexual generation of the *Neuroterus numismalis*and *Neuropterus quercusbaccarum* species, (i) button and (ii) spangle galls respectively, appear at the same time of the year on the same leaves. (e) button galls on an oak leaf, scale bar is 1 cm. (f) spangle galls on an oak leaf, scale bar is 1 cm. (g) CT scan of a button gall. (h) Zoomed in view of the button gall showing the trichomes. (i) CT cross section of the button gall, scale bar is 0.5 mm. (j) CT scan of a spangle gall. (k) Zoomed in view of the spangle gall showing the trichomes. (l) CT cross section of the spangle gall, scale bar is 0.55 mm.

Gall wasps do not induce uncontrolled cell division such as in *Agrobacterium* induced crown galls (Kerpen et al., 2019). Instead, they trigger a host-inducer species-specific developmental program for a new organ. The mechanism of induction is currently unknown. One hypothesis suggests that hormones such as auxins, cytokines and indole-3-acetic acid could be produced by the inducer to stimulate growth (Stone and Schönrogge, 2003). A second hypothesis is that secreted proteins could be involved in gall development. In aphids bicycle proteins have been shown to change plant gene expression and gall phenotype (Korgaonkar et al., 2021).

A third hypothesis is that galls may be induced through novel genes introduced via exogenous DNA insertions (Gatjens-Boniche, 2019, Cornell, 1983, Jankiewicz et al., 2017). It is known that parasitic Hymenoptera wasps have symbiotic viruses or virus like particles, which are injected with the egg into the host insect. These viruses help the developing larva to avoid the host immune system through gene transfer of virulence factors which disrupt capsule formation killing hemocytes, which are part of the insect immune system, or stopping them from adhering to foreign surfaces (Drezen et al., 2017, Moreau et al., 2009, Strand and Pech, 1995).

Currently viral particles have not been found in the ovaries or venom glands of two cynipid gall wasp species, *Biorhiza pallida* and *Diplolepis rosae* (Cambier et al., 2019, Hearn et al., 2019). However, this could be a limitation of the transcriptome annotations of these species, a high proportion of transcripts from the venom glands were novel transcripts and could be of non-wasp origin. Evidence of Wolbachia symbiotic bacteria was also found in *de novo* assemblies of both *B. pallida* and *D. rosae* (Hearn et al., 2019). In some non-cynipid systems, host manipulation by bacterial symbionts has been observed. For example, in leaf mining moths, successful gall formation requires the presence of Wolbachia (Nelson et al., 2014, Joy, 2013, Bansal et al., 2011, Kaiser et al., 2010). The Wolbachia bacteria modulates cytokine levels to create green islands in the senescent leaves where the galls are located, these islands provide a nutrient source for the developing larva (Kaiser et al., 2010). However, Wolbachia are not consistently found in gall wasps so are unlikely to play a fundamental role in gall formation (Rokas et al., 2002, Plantard et al., 1999).

The process of novel DNA insertions could be similar to crown galls, where a T-DNA insertion is required to hijack normal growth pathways to create a tumour like growth. This occurs by the T-strand integrating into the host plant genome and causing malignant growth though the expression of auxin, cytokinin and opine (Tiwari et al., 2022). Regardless of the mechanism, the complexity of the different gall forms suggests that a controlled and regulated re-programming of plant gene expression is taking place, which would be hard to achieve by simply modulating phytohormone levels.

Oak galls have specific external and internal structures, they are comprised of three different layers; nutritional, parenchymal, and epidermal. Each tissue layer provides a specific function, which differs significantly from the roles of the cells found in the original tissue. The galls attached to leaves have a broader impact on the leaf to which it is attached. Leaves with galls exhibit reduced photosynthesis, chlorophyll, and carotenoid levels (Kot et al., 2018b, Kot et al., 2020). Additionally, there are increases in free radicals and defensive responses (Kot et al., 2018a, Kot and Rubinowska, 2018, Kot et al., 2019).

Previous studies have compared gene expression in morphologically different galls formed on *Glochidion obovatum*, *Eurya japonica* and *Artemisia montana* which are induced by *Caloptilia cecidophora* a micromoth, *Borboryctis euryae* an aphid and *Rhopalomyia yomogicola* a gall midge respectively (Takeda et al., 2019). This group found that genes associated with photosynthesis decreased in all three gall types, and genes associated with “developmental processes” were upregulated, including 38 genes which are upregulated in all. The transcriptome of the oak gall generated by *B. pallida* on oak *Q. robur* has been sequenced, gene expression between galled and ungalled leaf tissue were markedly different. Expression of genes similar to Nod factor-induced early nodulin (ENOD) genes were expressed early in development (Hearn et al., 2019). Proteomic analysis has been used to investigate three gall types induced by *Cynips quercusfolii*, *Cynips longiventris*, and *Neuroterus quercusbaccarum*. This analysis identified 21 proteins that showed significant change in abundance in the galls but not in the host leaf. Many of these proteins were classed as involved in “developmental regulation of plant tissue into a gall” (Pawłowski et al., 2017). While the level of photosynthetic reduction and volatile production have been compared between spangle and button galls, the transcriptomes of these closely related but morphologically distinct galls have not been reported. Additionally, it is not known how the gene expression varies between the galls induced by the different asexual generations of the same species.

In this study, four galls from two wasp species were investigated. Button galls and spangle galls, containing the sexual generation and referred to here as sexual generation galls, are induced by *N. numismalis* and *N. quercusbaccarum* wasps respectively. Similarly, blister galls and currant galls, contain the asexual generation of *N. numismalis* and *N. quercusbaccarum* wasps respectively, and are referred to here as asexual generation galls. Previous studies have not sequenced or compared the asexual and sexual galls of the same species, or phenotypically distinct galls of closely related species that occur in the same ecological niche. Our study addresses these gaps by comparing the transcriptomes of the spangle, button, currant and blister galls. We expect that the galls that look more similar will have more similar gene expression profiles. To investigate the hypothesis that DNA from the inducer organism could integrate into the host to initiate oak galls, nanopore long read sequencing was conducted on the button and spangle galls. If insertions of wasp-derived DNA into the oak genome of initiator leaf cells were required for gall induction, we would expect to be able to detect these insertions flanked by known oak sequences.

## 2. Methods and Materials

### 2.1 Plant material

The four gall types were collected from the same *Quercus robur* tree. The sexual generation of *Neuroterus numismalis* (button gall) and *Neuroterus quercusbaccarum* (spangle gall) were collected in August 2020. Both button galls and spangle galls were pooled into groups of twenty to make up 50mg needed for DNA or RNA extraction. At the same time the leaf the galls were attached to was collected and leaves without any galls attached. Leaves were selected which only had one gall type on them. Major leaf veins were removed from the leaf samples in order to minimise the presence of different leaf structures, which could contribute RNA characteristic of a cell-type from which the galls were not derived. The asexual generation of *Neuroterus quercusbaccarum* (currant gall) was collected in May 2021, and the asexual generation of *Neuroterus numismalis* (blister gall) was collected in May 2022. These galls are larger, so 5 galls were pooled together to make up the 50mg needed for RNA extraction. At the same time the leaf the galls were attached to was collected. Galls and leaves were collected in triplicate. After collection galls were frozen in liquid nitrogen within 20 minutes and stored at −80 or processed straight away.

### 2.2 X-ray computed microtomography (CT) imaging and analysis

CT scans were carried out following a modified protocol by Lundgren et al. (Lundgren et al., 2019). Disks circa 5 mm in diameter were excised from the leaf encompassing the oak gall and then fixed to a polystyrene block on a plastic rod. Each sample was scanned using a GE phoenix nanotom X-ray µCT scanner (Waygate Technologies, Germany). Scan resolution was set at 2 µm, with an X-ray voltage of 72 kV and a current of 100 µA, collecting 2400 projections using a detector exposure time of 750 ms. Scan time was 30 minutes. 2D projections were reconstructed into 3D volumes using a filtered back-projection algorithm (Datos|X software, Waygate Technologies, Germany) prior to rendering and visualisation using VG StudioMAX software (Volume Graphics GmbH, Germany).

### 2.3 RNA extraction, library preparation, and RNA-Seq

RNA extraction was carried out on three biological replicates of the four galls (button, spangle, currant, blister) and the leaf that the galls were attached to. Each biological replicate consisted of 50mg of pooled gall or 50mg of leaf tissue the galls had been attached to. A modified CTAB method was used. (Pushkova et al., 2019, Barbier et al., 2019). 0.5ml of CTAB buffer (100mM Tris-HCL, pH9.5, 2% CTAB, 1.4M NaCl, 1% PEG 8000, 20mM EDTA, 2% PVP-40, 40mM DTT) and 2µl proteinase K was added to the ground tissue. This was incubated for 5 minutes at 65°C. 60µl of 10% SDS was then added, the tube inverted, and 1 volume of chloroform added. The samples were vortexed for 10 seconds, and centrifuged (4°C, 5500xg,10 minutes). The aqueous phase was taken and mixed with an equal volume of chloroform. After centrifugation (room temperature, 14,000xg, 5 minutes), the aqueous phase was taken and precipitated overnight with 0.5 volumes of 7.5M Ammonium acetate and 2.5 volumes of ethanol. After extraction the RNA was treated with turbo DNase (Invitrogen, AM2238) according to the manufacturer instructions.

Total RNA was sent to Novogene for poly-A enrichment library preparation and sequencing on the Illumina NovaSeq 6000 Sequencing System.

### 2.4 RNA-seq data processing

Low quality reads and adaptor sequences were removed using the default parameters of fastp v0.20.1 (Chen et al., 2018). Reads were then processed for compatibility with the transcripts based on pseudoalignment using the default parameters of kallisto v0.46.0 to quantify transcript abundances. For this quantification, the *Q. robur* transcriptome published by Plomion et al. was used (Plomion et al., 2018, Bray et al., 2016). Differential expression analysis was performed using sleuth using the default parameters (Pimentel et al., 2017). The differential expression compared the gene expression of the genes in the gall to those in the leaf they were attached to. Genes were classified as differentially expressed if they met the criteria of a q-value < 0.05 and log2 fold-change >= 1.5.

To annotate the differentially expressed genes with Ensembl IDs, to allow GO analysis to be performed, NCBI blastn was used to map the transcripts to those of the *Quercus lobata* transcriptome (VallyOak3.0, INSDC Assembly GCA_001633185.2). To annotate the differentially expressed genes with gene names this was repeated using the *Q. lobata* transcriptome from NCBI (GCF_001633185.2, version 3.2).

### 2.5 RNA-seq analysis and visualisation

To characterise the transcription factors that are differentially expressed in the four gall types, we compared a list of all known transcription factors in *Q. robur* with our list of DEGs (Zheng et al., 2016). To get the homologues of these genes for *Q. lobata* blastn was used to map the transcription factors to the *Quercus lobata* transcriptome (VallyOak3.0, INSDC Assembly GCA_001633185.2).

To determine enrichment the representation factor was calculated by an online tool from the Jim Lund lab, available at http://nemates.org/MA/. Gene Ontology (GO) functional enrichment of differentially expressed genes was conducted using Gprofiler with the organism *Q. lobata* (Raudvere et al., 2019). To visualise the data, venn diagrams were created using Deepvenn and the heatmap was generated using the R package heatmaply (Hulsen, 2022, Galili et al., 2017).

### 2.6 DNA extraction and nanopore sequencing

DNA was extracted from unaffected leaf, button gall and spangle gall. The button and spangle galls were pooled together, about 20 galls made up the 50mg of tissue. DNA extraction was performed using a modified CTAB (Cetyltrimethylammonium bromide) method (Pushkova et al., 2019, Barbier et al., 2019). 50mg of galls or leaf tissue was ground and combined with 1ml CTAB buffer (100 mM Tris-HCL, pH 9.5, 2% CTAB, 1.4M NaCl, 1% PEG 8000, 20mM EDTA, 2% PVP-40, 0.25% β-mercaptoethanol) and 30µl RNase A (New England Biolabs; T3018L). This was incubated for 40 minutes at 65°C before 10µl of proteinase K (New England Biolabs; P8107S) was added and incubated for a further 20 minutes. This was allowed to cool to room temperature before 120µl 10% SDS and 1 volume of chloroform was added. The samples were mixed by vortex for 10 seconds and centrifuged (4°C, 5,500xg, 10 minutes). The aqueous phase was taken and mixed with an equal volume of chloroform. After centrifugation (4°C, 5,500xg, 10 minutes), the aqueous phase was taken, and the DNA was precipitated overnight with 0.8 volumes of isopropanol at −20°C. After centrifugation (4°C, 5,500xg, 30 minutes), pellets were washed with 70% ethanol, dried, then resuspended in 50µl of water.

DNA for nanopore sequencing was further purified using a Qiagen genomic tip G/20 (Qiagen; 10223) following manufacturer instructions. The DNA was subsequently precipitated using Short Read Eliminator (SRE) XS kit (Circulomics; SS-100-121-01) to remove any fragments < 4 kb and progressively deplete fragments < 10 kb.

Sequencing libraries were prepared using the Genomic DNA Ligation Kit (Oxford Nanopore Technologies; SQK-LSK110) and approx. 2µg of SRE-XS treated DNA. 250ng of library was loaded on PromethION R9.4.1 flow cells (Oxford Nanopore Technologies; FLO-PRO002) on a PromethION Beta. During the sequencing runs, the Flow Cell Wash Kit (Oxford Nanopore Technologies; EXP-WSH004) was used halfway through to free up unavailable and blocked pores, after which the library was subsequently reloaded as before.

Base calling was performed using the Guppy high accuracy (HAC) base calling model 2.4.

### 2.7 Searching for genomic insertions

To search for genomic insertions in the galls a structural variant approach was taken. The nanopore sequencing from the button and spangle galls and unaffected leaf was assembled and then the gall and leaf genomes compared using structural variant analysis programs.

Raw nanopore reads were first examined via Nanoplot (v1.38.1) before being assembled and polished via flye (v2. 9) using default settings (De Coster and Rademakers, 2023, Lin et al., 2016). Each of these *de novo* assemblies were then assessed via BUSCO (v5.22) against the OrthoDB Odb10 eudictot database whilst contigs larger than 10,000 bp were further examined via QUAST (v5.02) using an upper bound assembly and settings for large, fragmented genomes (--min-contig 10000, -- upper-bound-assembly, --large, --fragmented) (Mikheenko et al., 2018, Kuznetsov et al., 2022, Manni et al., 2021).

Structural variants (SVs) were then examined independently via Assemblytics (v1.2.1), Minigraph (v0.15) and Sniffles (v1.0.12) (Nattestad and Schatz, 2016, Li et al., 2020, Sedlazeck et al., 2018). For Assemblytics, leaf tissue assemblies were first individually aligned against those of each gall tissues via MUMer (v3.23) using recommended parameters (-maxmatch, −l 100, −c 500) before calling SVs between 500 bp and 30,000 bp with no minimum unique sequence length (Kurtz et al., 2004). For Minigraph, SVs were called directly on alternative alignments performed between the same assemblies using long read mode (-x lr). For Sniffles, individual mappings were first performed for (i) leaf reads against the respective gall tissue assemblies as well as for (ii) the respective gall tissue reads against the leaf assembly via Minimap (v2.23) in nanopore mode and with MD tags included (-x map-ont, --MD) (Li, 2018). SVs were then called on the resulting sequence alignments using default settings and examined via SURVIVOR (v1.0.7) with the minimum read support disabled (Jeffares et al., 2017).

Putative indels between 500 bp and 30,000 bp were then extracted via custom python scripts and queried for any sequence similarities via MegaBLAST (v2.12). For this a custom database was used consisting of the haploid oak genome (PM1N), 32,393 microorganism genomes from the NCBI Refseq database with an assembly level of ‘chromosome’ or better (31,817 bacteria, 456 archaea, 79 fungi, 41 protozoa) in addition to 315 hymenoptera genomes from the NCBI Genbank database that were categorised as either ‘representative’ or ‘reference genome’. Amongst the latter, 22 assemblies belonged to members of the cynipidea family including *N. quercusbaccarum* (accession GCA900490065.1). Because no assembly was available for *N numismalis*, GCA900490065.1 also served as a proxy for this species.

In parallel, the respective nanopore reads were mapped to each of the *de novo* genome assemblies via Minimap. Those reads spanning the coordinates of putative indel sites that had matched with GCA900490065.1 were extracted via Samtools (v1.14) along with 5000 bp flanking sequences within the respective genome assemblies (Danecek et al., 2021). Extracted reads were then assessed via MegaBLAST using the same custom database and putatively described as either ‘oak’, ‘wasp’ or ‘chimeric’ according to whether they matched either and/or both PM1N and GCA900490065.1. These categorised reads were further filtered via custom python script to extract those which had matches for the respective genome assemblies longer than 1,000 bp and/or at least 80% percent identity.

## 3. Results

### 3.1 Gall characteristics and trichome morphology in Neuroterus species galls

We undertook CT scans; to better visualise the external and internal structures in live samples of the two sexual generation galls. These scans revealed the different layers of the gall and highlighted the rearrangement of trichomes on the gall surface compared to those on the leaf (Figure 1g-l).

Trichomes on *Q. robur* leaves are typically small and have a simple-uniseriate structure with enlarged basal cell and tapered apex (Jankiewicz et al., 2017). However, their morphology undergoes notable changes on the two gall types. On the button gall the trichomes change to a singular fasciculate structure characterised by long thick walled trichomes (Figure 1h). On the spangle gall trichomes have a more multiradiate structure which has thick walled trichomes which are fused at the base with the rays emanating at different levels (Figure 1k)(Hardin, 1976). These trichomes also significantly increase in size and number, and are a key feature used for gall classification.

CT imaging revealed other features, such as the attachment between the leaf and the gall is approximately 0.2mm in diameter. Additionally, a lighter line can be seen in the centre of both galls which is likely to be calcium oxalate crystals, a defence mechanism of the gall (Jankiewicz et al., 2021).

### 3.2 Comparative gene expression analysis in gall formation

To investigate altered plant biological process that are the same or different between the galls RNA sequencing was performed. Genes with significantly altered expression (± 1.5 logfold2 change, q-value < 0.05) relative to the host leaf were identified (Supplementary Data 1). To obtain gene names and Ensembl annotations for these genes, blastn was conducted against the *Q. lobata* transcriptome which is better annotated than *Q. robur* (Supplementary Table 1, Supplementary Data 2, 3).

There is a significant reprogramming of gene expression in all gall types, compared to the leaf they were attached to, with the most pronounced changes occurring in the galls of the sexual generations. In spangle and button galls 8458 and 9322 genes were upregulated, and 7063 and 6263 genes were downregulated, respectively (Supplementary Table 1). Notably, there is a substantial overlap between the gene expression profiles of spangle and button galls, where 48% of the differentially expressed genes (DEG) are common to both species. The representation factor for this overlap is 1.1 (p=9.5e-119) indicating a slightly larger overlap than expected (Figure 2a). In contrast, fewer genes were differentially expressed in galls of the asexual generation, with 4486 and 2567 upregulated and 4115 and 1327 downregulated genes identified for currant and blister galls, respectively (Supplementary Table 1). 18.2% of the DEG are shared between the two gall types, the representation factor is 1.5 (p=4.036e-118) suggesting a larger overlap than is expected (Figure 2b). The button and the blister galls share 12% of the DEGs with a representation factor of 0.9 (p=4.2e-18) and the spangle and currant galls share 22.4% of DEGs with a representation factor of 0.9 (p=3.7e-82) (Figure 2c, d). These observations indicate a slightly smaller overlap between the two gene sets than expected. When visualising the shared genes between the four gall types using a heatmap, it becomes evident that the two sexual generation galls, button and spangle, show more similarity to each other than to the asexual generation galls from the same species (Figure 2e).

**Figure 2.**
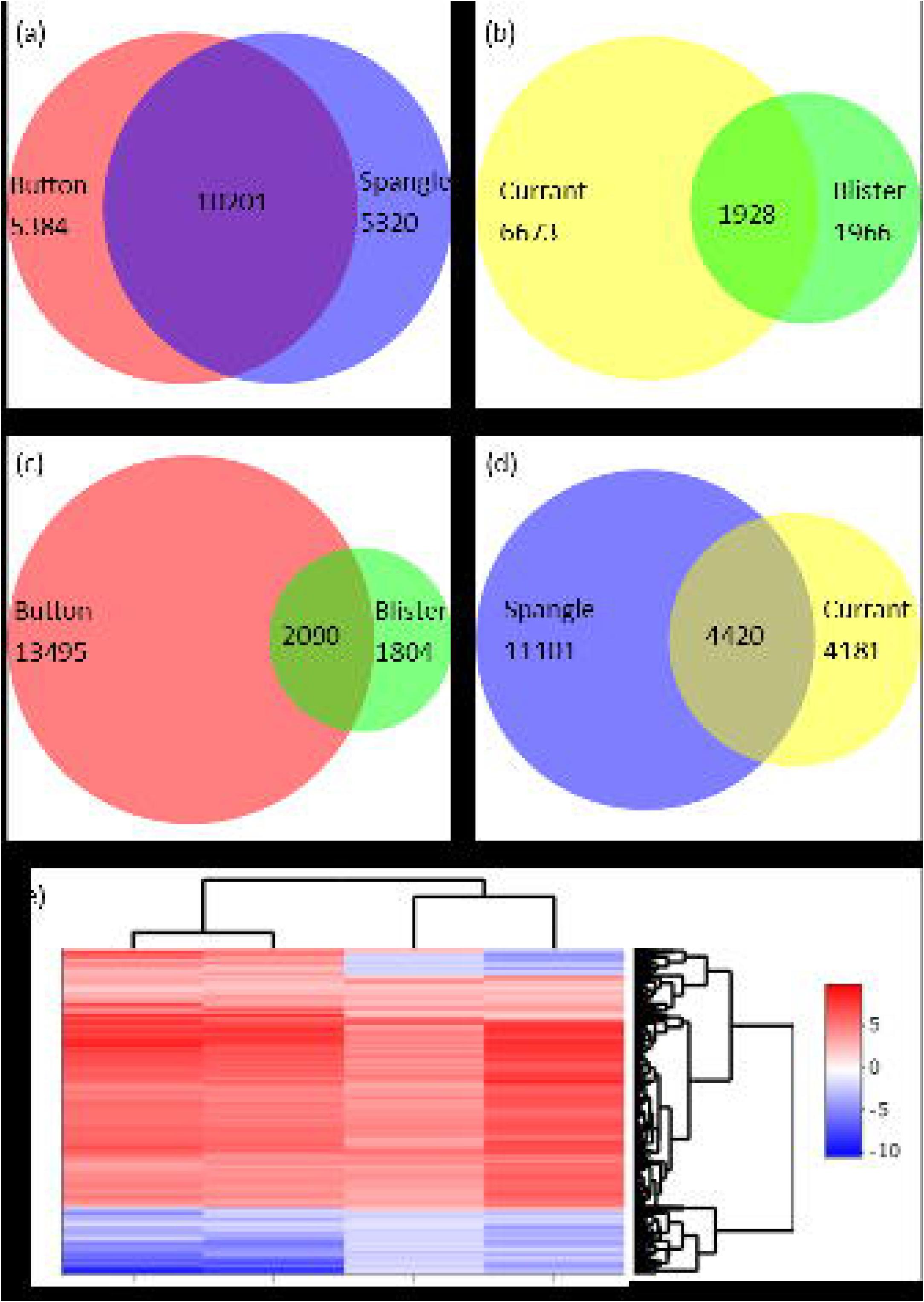
Significantly up and down regulated genes in the four galls. (a) Venn diagram of the significantly up- and down-regulated differentially expressed genes in the two sexual generation galls, button and spangle (qvalue 0.05, b +/- 1.5). (b) Venn diagram of the significantly up- and down-regulated differentially expressed genes in the two asexual generation galls, currant and blister (qvalue 0.05, b +/- 1.5). (c) Venn diagram of the significantly up- and down-regulated differentially expressed genes in the two galls from *N. numismalis,* button and blister (qvalue 0.05, b +/- 1.5). (d) Venn diagram of the significantly up and down regulated differentially expressed genes in the two galls from N*. quercusbaccarum*, spangle and currant (qvalue 0.05, b +/- 1.5). (e) Heat map of the 1251 significant differentially expressed genes found in all four galls.

Gene Ontology (GO) analysis was conducted on the DEGs that were shared between the galls in the comparison to identify overrepresented terms. These genes were broken down into upregulated, downregulated or where there was a mix of up- and down-regulated genes for the different galls in the comparison. When the four gall types are compared, terms such as ‘beta-fructofuranosidase’ and ‘fructose-bisphsphate aldolase activity’ are associated with the upregulated genes. An increase in energy demand through respiration correlates with the galls and their insect larva being an energy sink. In genes that change between the gall types, the terms ‘microtubule binding’ and ‘cytoskeleton motor activity’ are present. The genes associated with these terms are upregulated in the button and spangle galls and are downregulated in currant and blister galls. Interestingly, ‘fructose bisphophate aldolase activity’ reappears in the downregulated GO terms. When the genes associated with the GO term were examined, fructose bisphosphate 3 and 6 are upregulated and 1 is downregulated (Table 1).

**Table 1.** GO term analysis of the genes that are upregulated in all four galls, that change in expression between the four galls; and the genes that are downregulated in all four galls.

| GO term for upregulated genes |  |  |  |
| --- | --- | --- | --- |
| Source | Term ID | Term Name | Log qvalue |
| GO:MF | GO:0003824 | Catalytic activity | 2.529 |
| GO:MF | GO:0004564 | Beta-fructofuranosidase activity | 1.915 |
| GO:MF | GO:0008483 | Transaminase activity | 1.737 |
| GO:MF | GO:0004332 | Fructose-bisphosphate aldolase activity | 1.395 |
| GO:BP | GO:0005975 | Carbohydrate metabolic activity | 1.646 |
| GO term for genes that change between the galls |  |  |  |
| Source | Term ID | Term Name | Log qvalue |
| GO:MF | GO:0003774 | Cytoskeletal motor activity | 2.599 |
| GO:MF | GO:0022890 | Inorganic cation transmembrane transporter activity | 2.357 |
| GO:MF | GO:0008017 | Microtubule binding | 1.993 |
| GO:MF | GO:0016157 | Sucrose synthase activity | 1.494 |
| GO:BP | GO:0034220 | Monoatomic ion transmembrane transport | 2.054 |
| Go terms for downregulated genes |  |  |  |
| Source | Term ID | Term Name | Log qvalue |
| GO:MF | GO:0008194 | UDP-glycosyltransferase activity | 2.384 |
| GO:MF | GO:0004332 | Fructose-bisphosphate aldolase activity | 1.302 |

When the two asexual generation galls, currant and blister, are compared many terms are associated with the upregulated genes, including ‘response to an organic substance’ and ‘monocarboxylic acid binding’. The GO terms ‘nucleoside triphosphate diphosphatase activity’ and ‘nucleoside triphosphate catabolic process’ are upregulated in the blister gall but downregulated in the currant gall. ‘Phloem development’ is downregulated in both gall types (Table 2).

**Table 2.** GO term analysis of the genes that are upregulated, change between the galls, and are downregulated in the two asexual generation galls, blister and currant.

| GO terms for upregulated genes |  |  |  |
| --- | --- | --- | --- |
| Source | Term ID | Term Name | Log qvalue |
| GO:MF | GO:0016491 | oxidoreductase activity | 5.218 |
| GO:MF | GO:0019212 | phosphatase inhibitor activity | 3.838 |
| GO:MF | GO:0033293 | monocarboxylic acid binding | 3.290 |
| GO:MF | GO:0015144 | carbohydrate transmembrane transporter activity | 1.747 |
| GO:BP | GO:1901700 | response to oxygen-containing compound | 1.949 |
| GO:BP | GO:0010033 | response to organic substance | 1.811 |
| GO:BP | GO:1901575 | organic substance catabolic process | 1.509 |
| GO:BP | GO:0005975 | carbohydrate metabolic process | 1.303 |

**GO terms for genes that change between the galls**
| Source | Term ID | Term Name | Log qvalue |
| --- | --- | --- | --- |
| GO:MF | GO:0047429 | nucleoside triphosphate diphosphatase activity | 1.820 |
| GO:BP | GO:0009143 | nucleoside triphosphate catabolic process | 1.565 |

**GO terms for downregulated genes**
| Source | Term ID | Term Name | Log qvalue |
| --- | --- | --- | --- |
| GO:BP | GO:0010088 | phloem development | 3.898 |

When comparing the two sexual generations, spangle and button galls, several GO terms are associated with the upregulated genes, including ‘microtubule’, ‘nuclear division’ and ‘chromosome’. These GO terms suggest a high level of cell division is occurring. Some genes associated with the GO term ‘UDP glycosyltransferase’ are upregulated in the button gall and downregulated in the spangle gall, while other genes associated with this GO term are downregulated in both. Terms such as ‘photosynthesis’ and ‘thylakoid’ are downregulated in both galls, indicating that the galls are becoming energy sinks, taking energy from the leaf (Table 3).

**Table 3.**
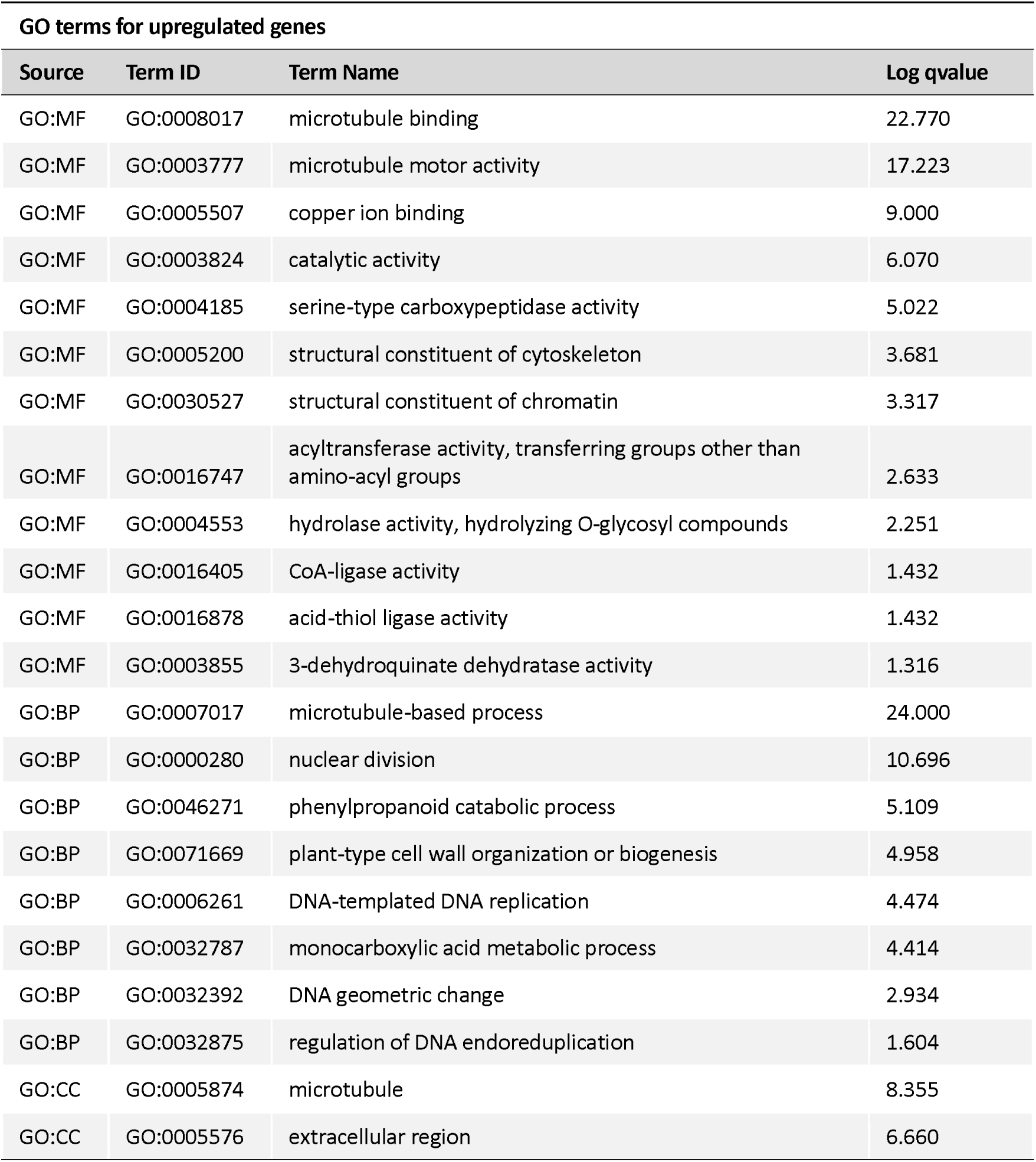
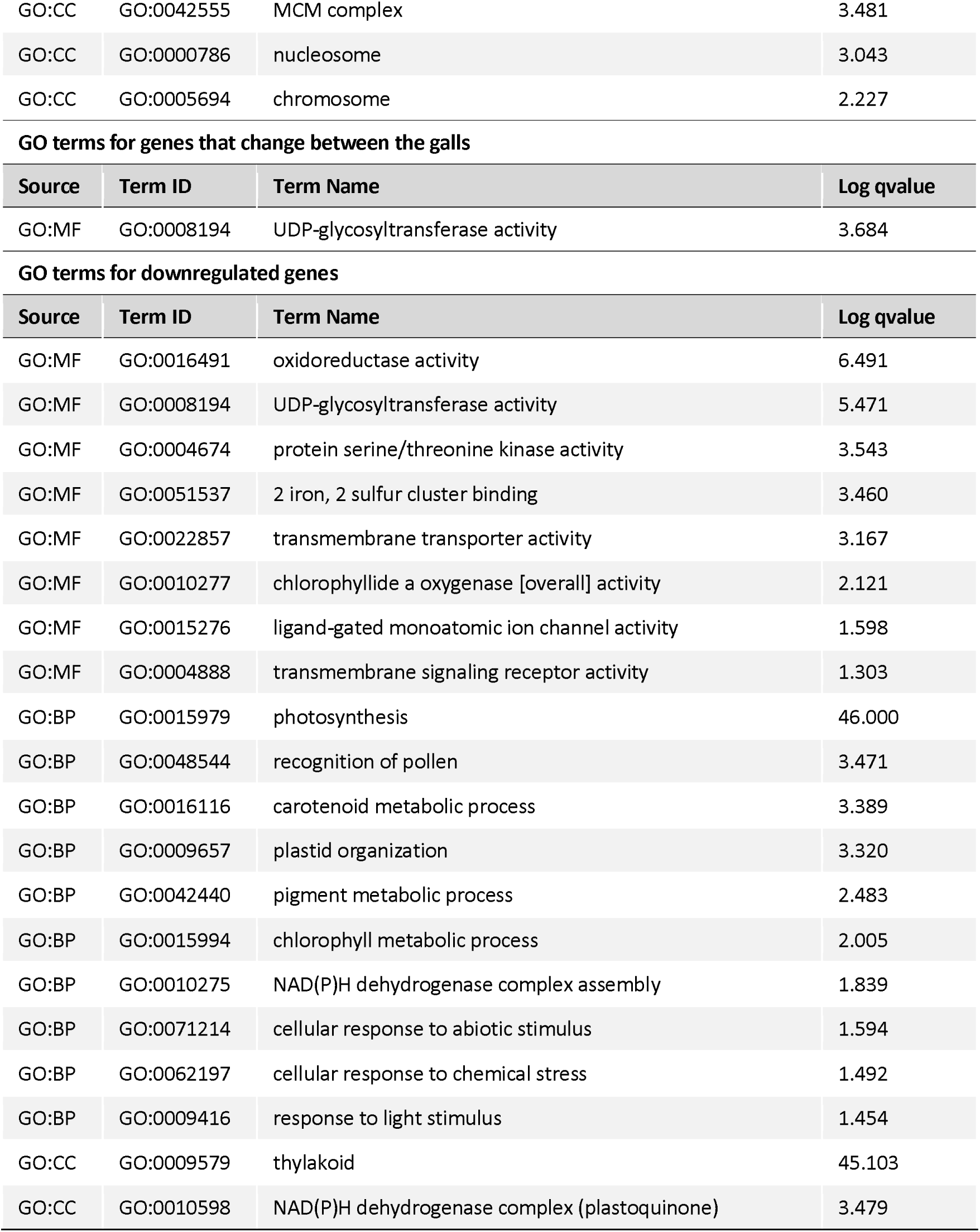
GO term analysis of the genes that are upregulated, change between the galls, and are downregulated in the two sexual generation galls, button and spangle.

**GO terms for upregulated genes**
| Source | Term ID | Term Name | Log qvalue |
| --- | --- | --- | --- |
| GO:MF | GO:0008017 | microtubule binding | 22.770 |
| GO:MF | GO:0003777 | microtubule motor activity | 17.223 |
| GO:MF | GO:0005507 | copper ion binding | 9.000 |
| GO:MF | GO:0003824 | catalytic activity | 6.070 |
| GO:MF | GO:0004185 | serine-type carboxypeptidase activity | 5.022 |
| GO:MF | GO:0005200 | structural constituent of cytoskeleton | 3.681 |
| GO:MF | GO:0030527 | structural constituent of chromatin | 3.317 |
| GO:MF | GO:0016747 | acyltransferase activity, transferring groups other than amino-acyl groups | 2.633 |
| GO:MF | GO:0004553 | hydrolase activity, hydrolyzing O-glycosyl compounds | 2.251 |
| GO:MF | GO:0016405 | CoA-ligase activity | 1.432 |
| GO:MF | GO:0016878 | acid-thiol ligase activity | 1.432 |
| GO:MF | GO:0003855 | 3-dehydroquinate dehydratase activity | 1.316 |
| GO:BP | GO:0007017 | microtubule-based process | 24.000 |
| GO:BP | GO:0000280 | nuclear division | 10.696 |
| GO:BP | GO:0046271 | phenylpropanoid catabolic process | 5.109 |
| GO:BP | GO:0071669 | plant-type cell wall organization or biogenesis | 4.958 |
| GO:BP | GO:0006261 | DNA-templated DNA replication | 4.474 |
| GO:BP | GO:0032787 | monocarboxylic acid metabolic process | 4.414 |
| GO:BP | GO:0032392 | DNA geometric change | 2.934 |
| GO:BP | GO:0032875 | regulation of DNA endoreduplication | 1.604 |
| GO:CC | GO:0005874 | microtubule | 8.355 |
| GO:CC | GO:0005576 | extracellular region | 6.660 |
| GO:CC | GO:0042555 | MCM complex | 3.481 |
| GO:CC | GO:0000786 | nucleosome | 3.043 |
| GO:CC | GO:0005694 | chromosome | 2.227 |

GO terms for genes that change between the galls
| Source | Term ID | Term Name | Log qvalue |
| --- | --- | --- | --- |
| GO:MF | GO:0008194 | UDP-glycosyltransferase activity | 3.684 |

GO terms for downregulated genes
| Source | Term ID | Term Name | Log qvalue |
| --- | --- | --- | --- |
| GO:MF | GO:0016491 | oxidoreductase activity | 6.491 |
| GO:MF | GO:0008194 | UDP-glycosyltransferase activity | 5.471 |
| GO:MF | GO:0004674 | protein serine/threonine kinase activity | 3.543 |
| GO:MF | GO:0051537 | 2 iron, 2 sulfur cluster binding | 3.460 |
| GO:MF | GO:0022857 | transmembrane transporter activity | 3.167 |
| GO:MF | GO:0010277 | chlorophyllide a oxygenase [overall] activity | 2.121 |
| GO:MF | GO:0015276 | ligand-gated monoatomic ion channel activity | 1.598 |
| GO:MF | GO:0004888 | transmembrane signaling receptor activity | 1.303 |
| GO:BP | GO:0015979 | photosynthesis | 46.000 |
| GO:BP | GO:0048544 | recognition of pollen | 3.471 |
| GO:BP | GO:0016116 | carotenoid metabolic process | 3.389 |
| GO:BP | GO:0009657 | plastid organization | 3.320 |
| GO:BP | GO:0042440 | pigment metabolic process | 2.483 |
| GO:BP | GO:0015994 | chlorophyll metabolic process | 2.005 |
| GO:BP | GO:0010275 | NAD(P)H dehydrogenase complex assembly | 1.839 |
| GO:BP | GO:0071214 | cellular response to abiotic stimulus | 1.594 |
| GO:BP | GO:0062197 | cellular response to chemical stress | 1.492 |
| GO:BP | GO:0009416 | response to light stimulus | 1.454 |
| GO:CC | GO:0009579 | thylakoid | 45.103 |
| GO:CC | GO:0010598 | NAD(P)H dehydrogenase complex (plastoquinone) | 3.479 |

When comparing the galls of *N. quercusbaccarum*, currant and spangle, genes associated with SAM (S-adenosylmethionine) are upregulated and GO terms such as ‘methionine adenosyltransferase activity’ and ‘S-adenosylmethionine biosynthetic process’ are present. Genes associated with the terms ‘DNA replication’ and ‘microtubule’ are upregulated in spangle galls and downregulated in currant galls. Common downregulated terms in both gall types include ‘photosynthesis’ and ‘thylakoid’ (Table 4).

**Table 4.** GO term analysis of the genes that are upregulated, change between the galls, and are downregulated in the two *N. quercusbaccarum* galls, currant and spangle.

GO terms for upregulated genes
| Source | Term ID | Term Name | Log qvalue |
| --- | --- | --- | --- |
| GO:MF | GO:0003824 | catalytic activity | 9.418 |
| GO:MF | GO:0004478 | methionine adenosyltransferase activity | 4.229 |
| GO:MF | GO:0003680 | minor groove of adenine-thymine-rich DNA binding | 1.951 |
| GO:MF | GO:0008483 | transaminase activity | 1.456 |
| GO:MF | GO:0030170 | pyridoxal phosphate binding | 1.449 |
| GO:BP | GO:0044281 | small molecule metabolic process | 4.917 |
| GO:BP | GO:0006556 | S-adenosylmethionine biosynthetic process | 3.964 |
| GO:BP | GO:0005975 | carbohydrate metabolic process | 3.833 |

GO terms for genes that change between the galls
| Source | Term ID | Term Name | Log qvalue |
| --- | --- | --- | --- |
| GO:MF | GO:0008017 | microtubule binding | 12.344 |
| GO:MF | GO:0003777 | microtubule motor activity | 10.610 |
| GO:MF | GO:0030527 | structural constituent of chromatin | 9.921 |
| GO:MF | GO:0052716 | hydroquinone:oxygen oxidoreductase activity | 6.288 |
| GO:MF | GO:0046982 | protein heterodimerization activity | 5.023 |
| GO:MF | GO:0005507 | copper ion binding | 4.699 |
| GO:BP | GO:0000280 | nuclear division | 13.491 |
| GO:BP | GO:0007018 | microtubule-based movement | 9.650 |
| GO:BP | GO:0046271 | phenylpropanoid catabolic process | 5.552 |
| GO:BP | GO:0071554 | cell wall organization or biogenesis | 2.796 |
| GO:BP | GO:0006260 | DNA replication | 2.669 |
| GO:BP | GO:0051172 | negative regulation of nitrogen compound metabolic process | 2.490 |
| GO:BP | GO:0005976 | polysaccharide metabolic process | 2.135 |
| GO:CC | GO:0000786 | nucleosome | 8.670 |
| GO:CC | GO:0048046 | apoplast | 4.918 |
| GO:CC | GO:0005874 | microtubule | 1.571 |

GO terms for downregulated genes
| Source | Term ID | Term Name | Log qvalue |
| --- | --- | --- | --- |
| GO:MF | GO:0005509 | calcium ion binding | 3.029 |
| GO:MF | GO:0016655 | oxidoreductase activity, acting on NAD(P)H, quinone or similar compound as acceptor | 2.238 |
| GO:MF | GO:0008194 | UDP-glycosyltransferase activity | 1.770 |
| GO:MF | GO:0004714 | transmembrane receptor protein tyrosine kinase activity | 1.666 |
| GO:MF | GO:0004888 | transmembrane signaling receptor activity | 1.615 |
| GO:BP | GO:0015979 | photosynthesis | 22.013 |
| GO:BP | GO:0010258 | NADH dehydrogenase complex (plastoquinone) assembly | 2.288 |
| GO:CC | GO:0009579 | thylakoid | 21.777 |
| GO:CC | GO:0010598 | NAD(P)H dehydrogenase complex (plastoquinone) | 5.974 |

When the *N. numismalis* galls, blister and button, are compared there are no GO terms associated with the genes that change between these two gall types. However, GO terms such as ‘Ent-kaurene oxidase activity’ and ‘ent-kaurene oxidation to kaurenoic acid’ are associated with the upregulated genes in both galls. This is interesting as ent-kaurene is an important step in gibberellin biosynthesis, a plant hormone that regulates many developmental processes including cell elongation.

Downregulated genes in both types of gall are associated with GO terms including ‘photosynthesis’ and ‘chloroplast’ (Table 5).

**Table 5.** GO term analysis of the genes that are upregulated, and are downregulated in the two *N. numismalis* galls, button and blister.

| GO terms for upregulated genes |  |  |  |
| --- | --- | --- | --- |
| Source | Term ID | Term Name | Log qvalue |
| GO:MF | GO:0052615 | ent-kaurene oxidase activity | 1.739 |
| GO:MF | GO:0003824 | catalytic activity | 1.333 |
| GO:BP | GO:0046394 | carboxylic acid biosynthetic process | 2.041 |
| GO:BP | GO:0009058 | biosynthetic process | 1.776 |
| GO:BP | GO:0010241 | ent-kaurene oxidation to kaurenoic acid | 1.317 |
| GO terms for downregulated genes |  |  |  |
| Source | Term ID | Term Name | Log qvalue |
| GO:MF | GO:0008194 | UDP-glycosyltransferase activity | 4.659 |
| GO:BP | GO:0009765 | photosynthesis, light harvesting | 1.467 |
| GO:CC | GO:0009507 | chloroplast | 3.440 |

### 3.3 Transcription factor expression and trichome development in gall formation

In the majority of the transcription factor families there is a range in the fold change of the genes. Fewer transcription factors are differentially expressed in the blister gall, with 104 identified, compared to the other three where there are 310 in the button gall, 286 in the spangle gall, and 202 in the currant gall (Supplemental Data 4). This observation is consistent with the appearance of the blister gall, which is more similar to the leaf than are the other galls (Figure 3).

**Figure 3.**
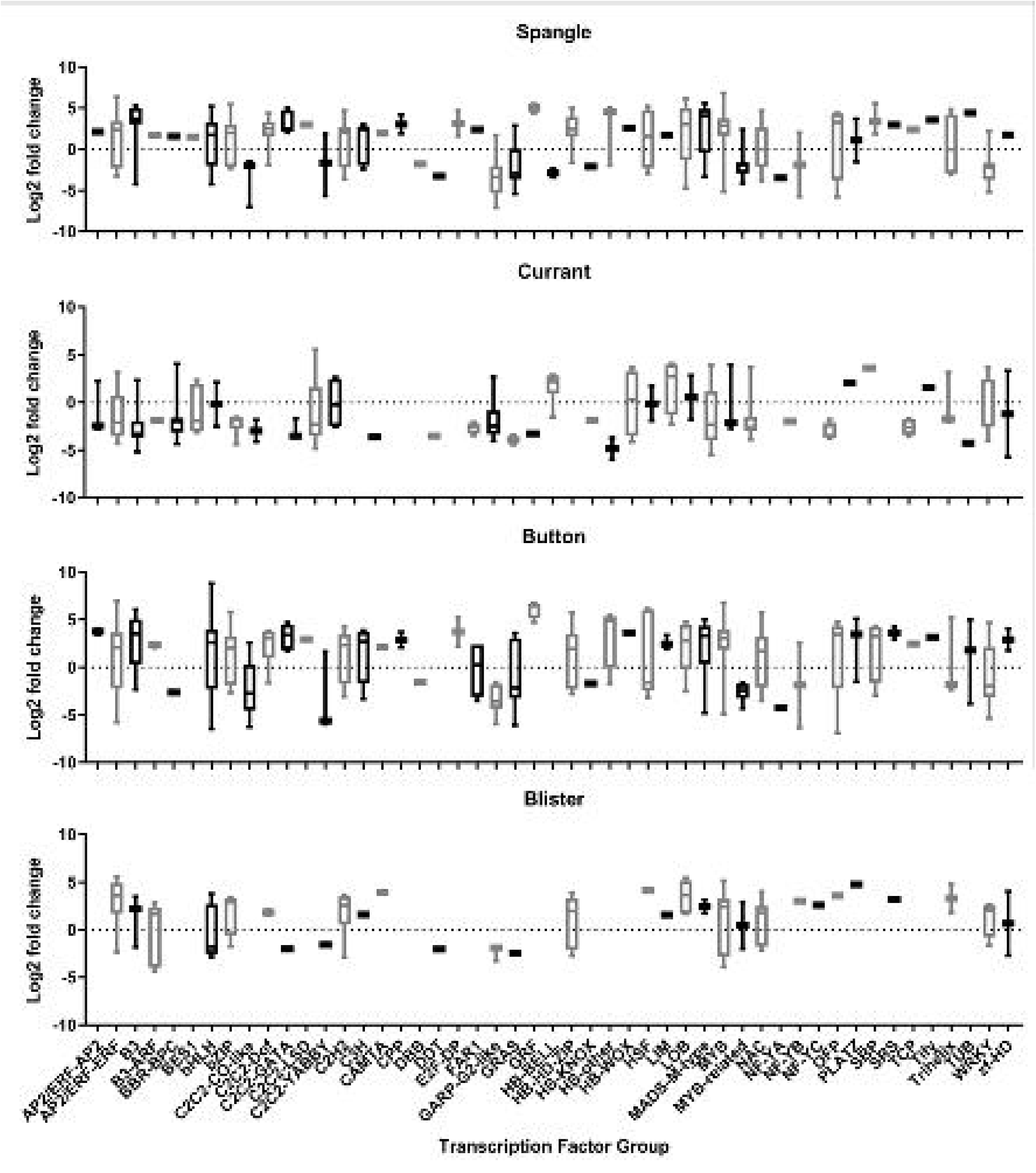
Fold change in transcription factor groups in the four gall types

There is a noticeable alteration in trichomes between the button and spangle galls and oak leaves. Button galls have a singular fasciculate form at a high density, while spangle galls have a multiradiate form. The development of trichomes is controlled by a complex network of positive and negative regulators. Among these regulators, ENHANCER OF GLABRA3 (EGL3) is part of the activator complex for trichome development (Payne et al., 2000, Zhang et al., 2003). In button and spangle galls this gene was observed to be upregulated with a 2.4 log2 fold change (q=6.98e-12) and 2.9 log2 fold change (q=3.14e-8) respectively. EGL3 it is not differentially expressed in the currant or blister galls, which lack trichomes.

### 3.4 There is no evidence of novel DNA insertions in the oak genome of sexual generation galls

It has been hypothesised that DNA insertions into the host genome could trigger the development of the gall structure, in a mechanism potentially analogous to the induction of crown gall tumours by *Agrobacterium tumefaciens* (Chilton et al., 1980). To explore this hypothesis, we conducted Nanopore long read sequencing of genomic DNA from button and spangle galls, as well as from a leaf from the host plant which did not have any galls attached to it. 86 GB of Oxford Nanopore Technologies (ONT) data for leaf (consisting of 7,198,195 reads with a mean read length of 11,954.7 and mean Q score of 13); 96 GB of ONT data for button gall (consisting of 5,284,520 reads with a mean read length of 18,277.7 and a mean Q score of 12.9) and 50 GB of ONT data for spangle gall (consisting of 5,116,190 reads with a mean read length of 9961 and a mean Q score of 12.5) were assembled using Flye. This resulted in a leaf genome with a N50 of 255784 and a length of 1175651838 bp; button genome with a N50 of 317134 and a length of 1697337048; a spangle genome with a N50 of 173653 and a length of 1269969265 (Table 6). A structural variant approach was taken to find potential insertions in the gall genomes, by comparing the spangle or button gall genome with the leaf genome to see if insertions or deletions could be found between them.

**Table:6.**
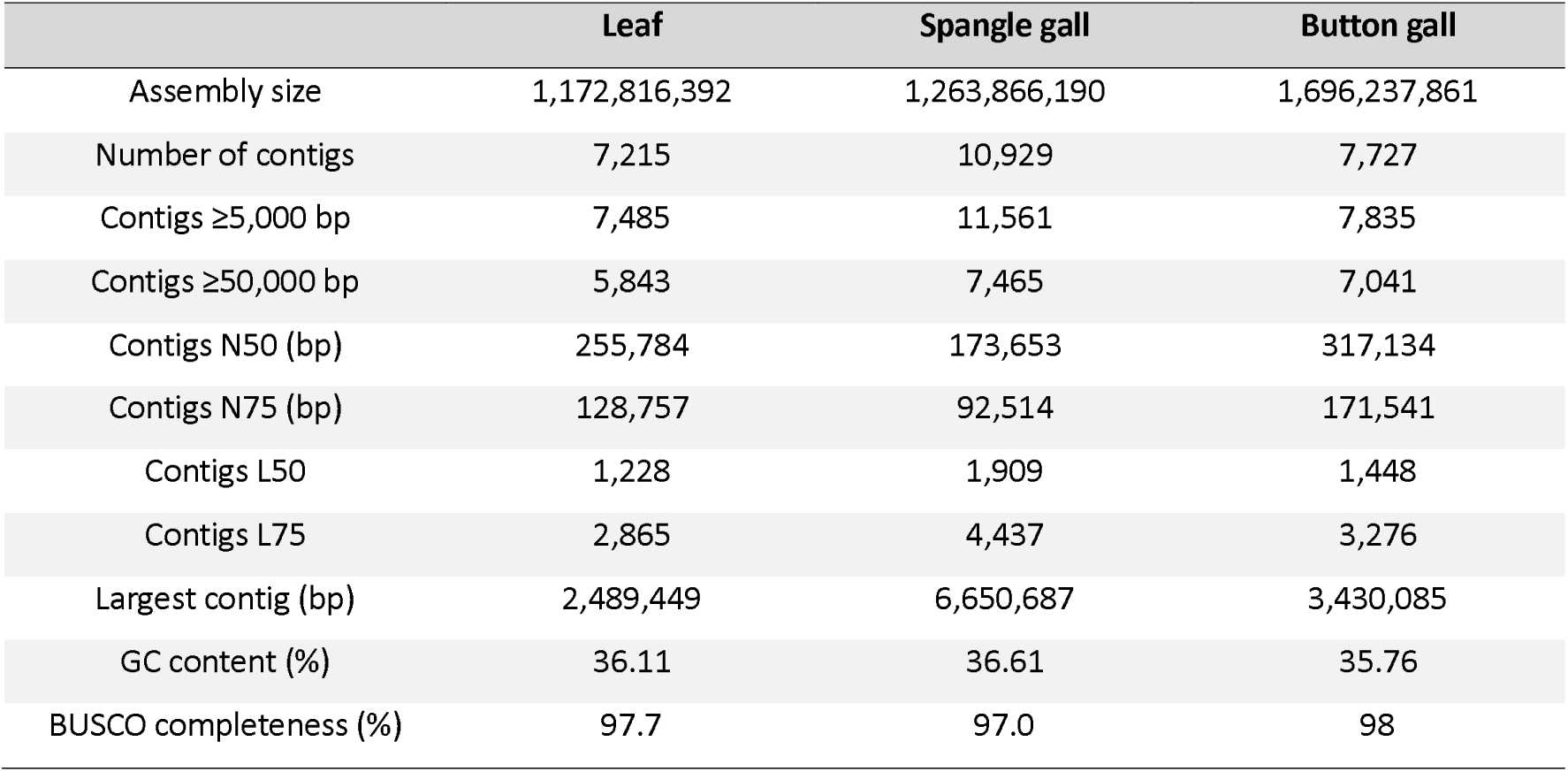
Summary of the genome assemblies.

Through the application of three independent structural variant approaches, thousands of potential insertions in the button and spangle galls were identified. The potential insertions were assessed to see if they matched the oak genome or the wasp, *N. querousbaccarum,* genome by mega blast, *N. numismalis* is not currently sequenced. Insertions that matched the respective genome assemblies for longer than 1000bp and/or at least 80% identity were further analysed (Supplemental data 5). Subsequent examination of the putative insertions revealed that they did not resemble any known DNA insertion mechanism. The putative insertions more closely resemble and were found in ribosomal gene clusters and spacer sequences. However, it remains unclear whether these findings are artefacts of the assembly process or expansions in these regions.

## 4. Discussion

### 4.1 Comparative imaging of the sexual generations

This study aimed to investigate the similarities and differences between the asexual and sexual generations of *N. quercusbaccarum* and *N. numismalis*. The CT imaging of the two sexual galls showed distinct differences in trichome organization. The trichomes on the spangle gall have a multiradiate structure, whilst on the button gall, they have a solitary fasciculate structure. The trichomes on both gall types are much larger and at higher density than the ones found on the host leaf.

The CT imaging also revealed the small size of the attachment, approximately 0.2 mm, and showed the previously described calcium oxalate layer surrounding the larval growth chamber (Jankiewicz et al., 2021). The CT imaging shows the density differences of the internal structures of the gall and enables a 3D image of the entire gall structure to be built. This approach provides a different prospective to the scanning electron images of these two galls currently published (Jankiewicz et al., 2017, Jankiewicz et al., 2021).

### 4.2 Novel insights into oak gall transcriptomes and transcription factor regulation

The gall structures have a very different appearance to the leaf to which they are attached, and so it is implicit that the transcriptomes of the leaf would be very different to that of the galls. The transcriptomes of several oak galls induced by various species of gall wasp have been reported (Hearn et al., 2019, Martinson et al., 2022). However, this study is the first analysis of *N. quercusbaccarum* and *N. numismalis* gall transcriptomes, and to our knowledge, the first comparison between asexual and sexual gall generations. Sequencing of the succulent oak gall induced by *Dryocosmus quercuspalustri s*on *Quercus rubra* revealed general trends, such as a decrease in photosynthesis and an increase in glycolysis (Martinson et al., 2022), similar to the gall types analysed here. In oak apples, which are induced on *Q. robur* by *B. pallida,* gene expression patterns diverge from normal bud development as the gall matures (Hearn et al., 2019).

We have revealed that galls which are found in the same niche at the same time have similar global reprogramming of plant gene expression. Button and spangle galls have a large overlap between their transcriptional profiles, with 48% of their DEGs being shared. This observation may be attributed to several factors; both galls are induced on the same leaf stage, and leaf stage has been documented to have a crucial role in successful gall initiation (Hough, 1953). Additionally, the larva hibernates inside the gall over winter, these similar internal structures imply similar gene expression.

The observed changes in transcription factor expression likely underpins the global transcriptional changes that result in formation of galls with wasp-specific phenotypes. To better understand the gall forming process, it is necessary to unravel the hierarchy, regulatory networks, and mechanism(s) of initial induction of these transcription factors, as well as their target genes. There are 1163 transcription factors in the ITAK list for *Q. robur* (Zheng et al., 2016). In the button and spangle galls a higher number of transcription factors were differentially expressed, 310 and 286 respectively, compared to the currant and blister galls 202 and 104 respectively. One transcription factor, EGL3, a gene known to be required for trichome initiation is highly expressed in button and spangle galls, though not differentially expressed in the galls of the trichomeless asexual generation. Transcription factor expression in oak galls of other species has not been specifically investigated. However, in wild grape vines (*Vitis riparia*), the expression of MYB33 is downregulated in late stage galls induced by the phylloxera *Daktulosphaira vitifoliae*, suggesting a role in supressing gibberelin signalling and potentially floral development in these structures (Schultz et al., 2019).

### 4.3 Potential mechanisms for parasitoid-induced gall formation

Our analysis detected thousands of structural variants in the genomes of the spangle and button gall. However, the sequences of the structural variants that could be found in both *N. quercusbaccarum* and the respective gall genome were ribosomal sequences. These potential rearrangements or insertions are found in areas which conserved throughout the evolutionary tree and occur at many loci, and so simply may be due to local repeat expansions or contractions. The genome assemblies that we produced for this work have a high completeness and however the contiguity is low but comparable to other tree genomes (Plomion et al., 2018).

While the ribosome was long thought of as just the factory that made the proteins required for the cell, there is increasing evidence to support its role in translation regulation (Genuth and Barna, 2018). Ribosomal heterogeneity has been linked to key roles in development and differentiation in many organisms from humans to plants (Norris et al., 2021, Martinez-Seidel et al., 2020). Ribosomal expansions have been demonstrated to play important functional roles as well as providing protein binding sites (Fujii et al., 2018). Ribosomal proteins have also been linked to galling diseases, such as clubroot, where changes in ribosomal proteins are used as markers to detect the geographic origin of the strain as well as its virulence (Laila et al., 2017, Javed et al., 2023). Ribosomal proteins in rice have also been linked to resistance against gall midges (Moin et al., 2021).

The results from this study provide an insight into the altered gene expression associated with asexual and sexual galls produced by two closely related wasp species, *N. quercusbaccarum* and *N. numismalis.* We did not find evidence of foreign DNA insertions into the oak genome of the gall, although there was some evidence of structural rearrangements (rRNA genes) of the genome. It remains an interesting question how a few cells in the leaf are induced to divide and differentiate into such structurally defined and characteristic gall forms.

## Acknowledgments

This work was supported by the Leverhulme Trust, Grant/Award Number; RPG-2020-284. We acknowledge the following people for their contribution to the study. Nanopore sequencing was conducted by Deepseq (University of Nottingham). The CT scanning was performed at the Houndsfield facility (University of Nottingham) by Dr Craig Sturrock. We also thank Laura Holt for her photography of the oak galls.

## Data availability

The RNA sequencing data reported in this paper has been deposited in the NCBI GEO database, accession number. GSE244168 (https://www.ncbi.nlm.nih.gov/geo/query/acc.cgi?acc=GSE244168). The DNA sequencing reported in this paper has been deposited in the SRA database Bioproject ID number. PRJNA1027974 (https://www.ncbi.nlm.nih.gov/sra/PRJNA1027974).

