## Supplementary table 2 for "Comparative transcriptome reprogramming in oak galls containing asexual or sexual generations of gall wasps"

| **Program** | **Comparison** | **Count** |
| --- | --- | --- |
| Assemblytics v1.2.1 | Leaf vs spangle gall | 3110 |
| Assemblytics v1.2.1 | Leaf vs button gall | 4089 |
| Minigraph v0.15 | Leaf vs spangle gall | 1341 |
| Minigraph v0.15 | Leaf vs button gall | 935 |
| Sniffles v1.0.12 | Spangle gall vs leaf | 5908 |
| Sniffles v1.0.12 | Leaf vs spangle gall | 3836 |
| Sniffles v1.0.12 | Button gall vs leaf | 3637 |
| Sniffles v1.0.12 | Leaf vs button gall | 5440 |

Supplemental table 2. The number of possible insertions from the structural variant analysis comparing the gall genomes to the leaf control genome. Three programs were used assemblytics v1.2.1, minigraph v0.15 and sniffles v1.0.12 to detect insertions that were between 500 and 30,000 base pairs long.
