## Supplementary table 1 for "Comparative transcriptome reprogramming in oak galls containing asexual or sexual generations of gall wasps"

|  | Currant gall vs attached leaf | | | Spangle gall vs attached leaf | | | Blister gall vs attached leaf | | | Button gall vs attached leaf | | | |
| --- | --- | --- | --- | --- | --- | --- | --- | --- | --- | --- | --- | --- | --- |
|  | Transcript | Gene name | Ensembl ID | Transcript | Gene name | Ensembl ID | Transcript | Gene name | Ensembl ID | | Transcript | Gene name | Ensembl ID |
| Total up | 4486 |  |  | 8458 |  |  | 2567 |  |  | | 9322 |  |  |
| *Quercus lobata* |  | 2485 | 2518 |  | 6473 | 6567 |  | 1714 | 1717 | |  | 7020 | 7147 |
| Total down | 4115 |  |  | 7063 |  |  | 1327 |  |  | | 6263 |  |  |
| *Quercus lobata* |  | 3779 | 3837 |  | 6428 | 6543 |  | 1198 | 1202 | |  | 5832 | 5919 |

Supplementary table 1. The transcriptome that was used in the alignments was not annotated. To get annotations the sequences were nucleotide blasted against *Quercus lobata* to get the gene names and ensemble gene IDs. The table contains the number of significant up and down regulated genes (qvalue 0.05, fold change +/-1.5) for the differential expression of the gall types as well as how many results came back in the blast.
